# Development of a Consensus Molecular Classifier for Pancreatic Ductal Adenocarcinoma

**DOI:** 10.1101/2025.03.06.641837

**Authors:** Pablo Villoslada-Blanco, Lola Alonso, Sergio Sabroso-Lasa, Miguel Maquedano, Lidia Estudillo, Francisco X Real, Evangelina López de Maturana, Núria Malats

## Abstract

Pancreatic ductal adenocarcinoma (PDAC) presents a significant challenge, with a five-year survival rate of approximately 10%. Tumor heterogeneity contributes to the limited effectiveness of treatments. Several tumor and stroma molecular classifiers have attempted to clarify this heterogeneity with moderate agreement. Recognizing the complexity introduced by this extensive array of taxonomies, this study aims to develop a consensus molecular classifier by including both tumor and stroma features. We integrated gene expression data through Virtual Microdissection and classified the training samples to apply Machine Learning algorithms for each previous classifier. The consensus classifier was then derived using a Markov Clustering Algorithm, and its association with overall survival was assessed. The results indicated that Elastic-Net emerged as the superior model. We identified two classes for tumor components (Consensus Classical and Consensus Non-classical) and stroma components (Consensus Normal-Immune and Consensus Activated-ECM). The consensus Random Forest achieved a balanced accuracy of 96.33% and 98.92%, respectively. While not consistent across retrospective series, the algorithm (*PDAConsensus*) independently predicted overall survival. We developed a robust consensus classifier for PDAC that integrates tumor and stroma features and made it accessible through the R package *PDACMOC* (*PDACMolecularOmniClassifier*, https://github.com/pavillos/PDACMOC) and a Shiny app (https://pdacmoc.cnio.es/).

## INTRODUCTION

Pancreatic Ductal Adenocarcinoma (PDAC) is the predominant malignant tumor of the pancreas and the third leading cause of cancer death (1). It stands as one of the deadliest cancers, with a five-year survival rate of around 10% (2). Conventional treatment strategies have not demonstrated significant benefits for overall PDAC patients. This fact is partially attributed to PDAC heterogeneity and the potential variations in treatment responses among distinct molecular subtypes within the same cancer type. This observation has redirected recent research toward deciphering diverse transcriptional categorizations (3). Instead of relying solely on morphological attributes, molecular cancer classifications pivot towards the gene expression profiles of tumors. These classifications can spotlight specific gene attributes, such as mutations or varied expressions, offering a more nuanced understanding of the disease (4) and supporting research and patient clinical management.

On the other hand, it is worth mentioning that PDAC tumors possess a uniquely dense stromal environment, also known as the tumor microenvironment (TME). This stroma comprises a complex mix of fibroblasts, immune cells, vasculature, and extracellular matrix (ECM). Notably, the dense stromal component can account for up to 90% of the tumor volume in PDAC, presenting substantial challenges for treatment (5,6). This stroma not only acts as a physical barrier, preventing chemotherapeutic agents from effectively reaching the tumor cells, but also establishes a microenvironment that supports tumor growth, invasion, and resistance to therapy (7). Furthermore, the crosstalk between the stroma and cancer cells is believed to play pivotal roles in progression and metastasis (5,8). Recent studies have also elucidated the existence of distinct stroma subtypes within the PDAC microenvironment, which can have differential effects on tumor progression, therapy resistance, and patient outcomes (7,9), and, therefore, should be also considered as potential therapeutic targets.

PDAC tumor molecular classifiers have evolved significantly, with several key studies identifying distinct tumor subtypes based on gene expression profiles, each associated with unique clinical outcomes and therapeutic responses. Collisson et al. (10) identified three subtypes: classical, quasi-mesenchymal (QM), and exocrine-like, using transcriptional profiles and a 62-gene signature called PDAssigner. Moffitt et al. (11) differentiated tumor, stromal, and normal cell expressions through virtual microdissection (VM), revealing classical and basal-like subtypes, with the basal-like showing worse outcomes. Bailey et al. (3) reported four subtypes: squamous, pancreatic progenitor, immunogenic, and aberrantly differentiated endocrine-exocrine (ADEX), highlighting distinct prognoses and histopathological characteristics. Puleo et al. (12) combined stromal and neoplastic components into a new classification, identifying subtypes like pure classical, immune classical, pure basal-like, stroma activated, and desmoplastic, emphasizing the role of the tumor microenvironment in prognosis. Chan-Seng-Yue et al. (13) further classified tumors into five subtypes: classical-A, classical-B, basal-like-A, basal-like-B, and hybrid, noting differences in clinical stage and intratumoral heterogeneity.

On the other hand, recent studies on stroma molecular classifiers have shed light on the stroma component’s significant role in PDAC prognosis. Moffitt et al. (11) identified normal and activated stroma subtypes, highlighting the poor prognosis associated with the combination of activated stroma and basal tumor subtypes. The normal stroma subtype features higher expression of markers for pancreatic stellate cells, whereas the activated stroma subtype shows an upregulation of genes linked to macrophages. Maurer et al. (14) further delved into the interaction between tumor epithelial and stroma compartments, identifying immune-rich and ECM-rich stroma subtypes. The immune-rich subtype is noted for its immune signaling, while the ECM-rich subtype, associated with extracellular matrix pathways, correlates with lower survival rates.

Such a diverse range of classifiers poses challenges in identifying the most suitable one for research and clinical use. Besides, the intratumoral heterogeneity, the different processing methods, and the overlapping among the different subtypes, as highlighted by Collisson et al. (15), make classifying tumors even more difficult. Therefore, we aimed to generate a consensus molecular classifier integrating the information of all the previous classifiers, acknowledging other authors who have implemented such approaches in other cancers, such as colorectal (16) and bladder cancer (17).

## MATERIAL AND METHODS

### Study Population, Expression Data Retrieval & Preprocessing

We used mRNA expression data of 514 PDAC samples from The Cancer Genome Atlas (TCGA) (https://www.cancer.gov/ccg/research/genome-sequencing/tcga) (n=149), the International Cancer Genome Consortium (ICGC) (https://www.icgc-argo.org/) Canada (n=68 from batch 1 (ICGC_v100), and n=190 from batch 2 (ICGC_v84)), and the PanGenEU study (18,19) (n=107) to develop the consensus classification system. Two additional studies, ICGC Australia (ICGC-au, n=86) and an additional PanGenEU data (PanGenEU-validation set (PanGenEU-vs), n=67), were used as validation sets. Demographic and clinicopathological information were available for all datasets except for ICGC_v100.

Common genes for all datasets were selected, and a batch effect correction was applied using the ComBat_seq function from the sva R package (20). Then, low-expressed genes (genes with fewer than five reads in 50% of the samples) were filtered out, and a Variance-Stabilizing Transformation (VST) was applied using the varianceStabilizingTransformation function from the DESeq2 R package (21). This was followed by scaling using the scale function from R base.

### Tumor and Stroma Single Sample Classifiers

To build the tumor single sample classifiers (SSCs), we first considered the gene signatures defining each classification system to classify the tumors according to each one: Collisson’s included 56 genes (out of the 62 genes included in Figure 1 of Collisson et al. (10)); Moffitt’s consisted of 49 genes (from the Figure 3 of Moffitt et al. (11)), Bailey’s contained 251 genes out of the 100 top genes by log2FoldChange (log2FC) for each signature from the Significance Analysis of Microarray included in the Supplementary Table 14 of Bailey et al. (3); and Chan-Seng-Yue’s included 372 genes of the top 100 genes of each signature in Supplementary Table 4 and Figure 2b of Chan-Seng-Yue et al. (13). Then, separately for each classifier, we estimated the differential expression of each gene (log2FC) in the tumors assigned to a cluster compared to the rest using the log2FC function from the singleseqgset R package. Finally, we performed a gene set enrichment analysis (GSEA) over these log2FC using fgsea R package by comparing each cluster against the list of the gene signatures, allowing us to assign each cluster to a subtype.

**Figure 1.**
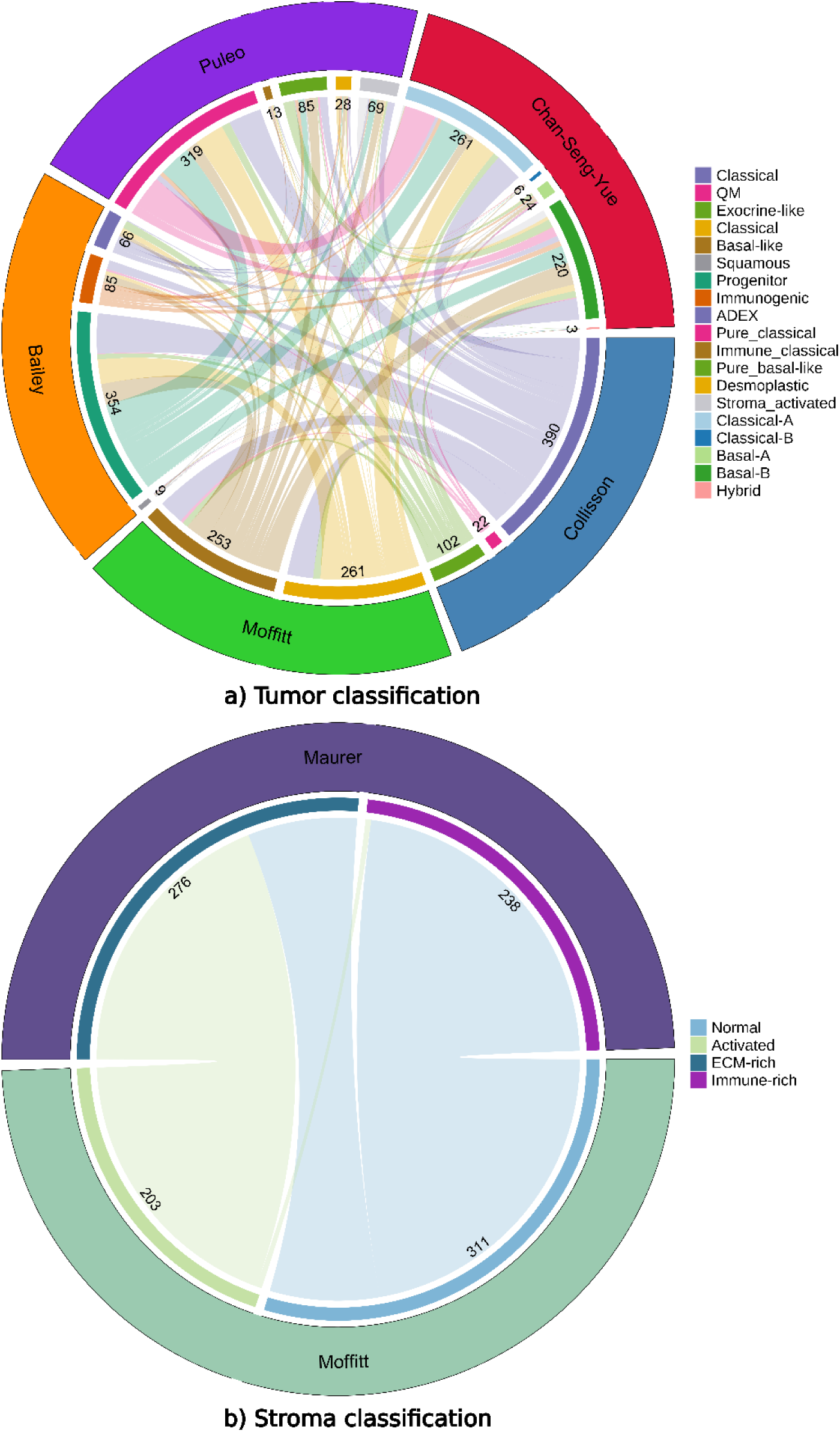
Circus plots show the distribution of the subtypes across the different classification systems. ADEX (aberrantly differentiated endocrine-exocrine), QM (quasi-mesenchymal).

**Figure 2.**
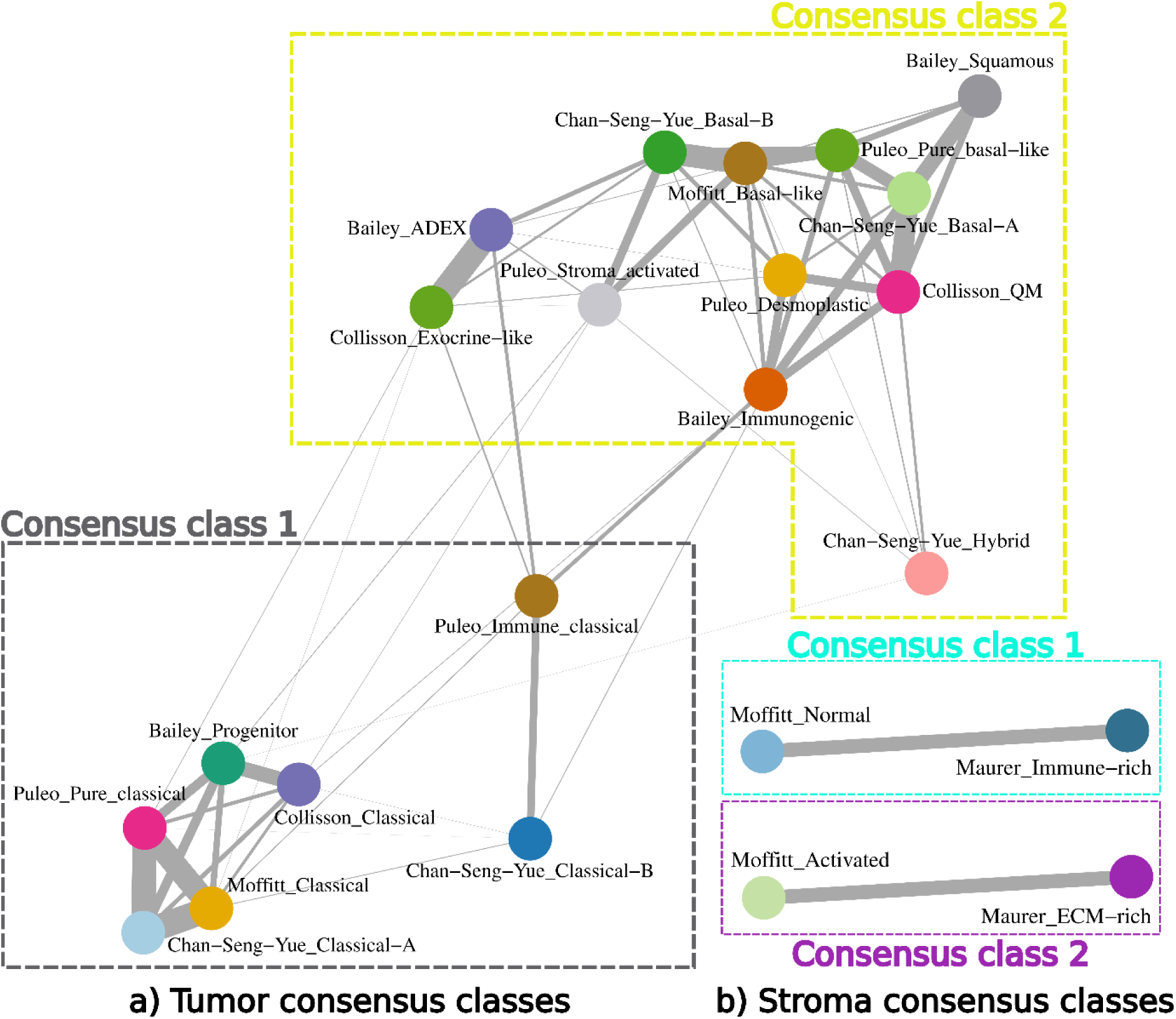
Consensus classes are derived from applying Markov Clustering Algorithm over the network of Cohen’s kappa indices between the subtypes of each classification.

**Figure 3.**
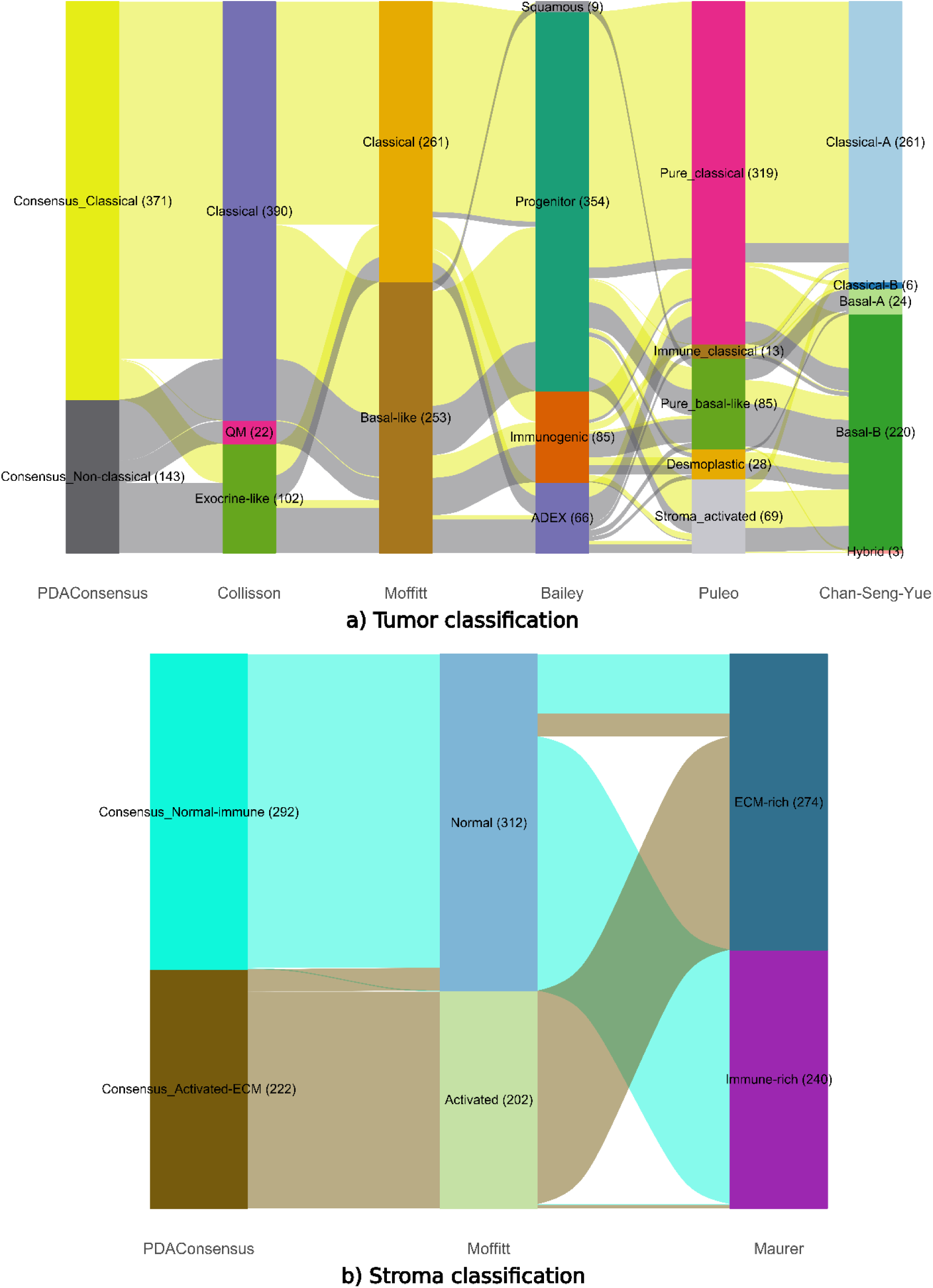
Parallel sets illustrate how samples classified as Consensus Classical or Consensus Non-classical and as Consensus Normal-immune or Consensus Activated-ECM are categorized by other classifiers.

Since no clear gene signatures are available to classify the tumors according to Puleo’s system, we first considered the expression of the genes defining the centroids provided in Supplementary Table 2 of Puleo et al. (12) (385 out of the 403 genes). Then, we calculated the Pearson’s correlation (*r*) between each sample and the centroids defining each subtype and assigned each sample to the subtype whose centroid showed the highest correlation coefficient (*r*) (cor function from stat R package).

We also classified the tumor samples according to the available stroma classifications. We first obtained the stroma compartment of each sample through VM using the ADVOCATE R package (https://github.com/califano-lab/ADVOCATE) (22). These gene signatures were composed of 43 genes in the case of Moffitt (out of the 48 genes in Figure 2 of Moffitt et al. (11)) and 468 genes in the case of Maurer (out of the 472 genes in Table S19 of Maurer et al. (14)).

We then trained a set of Machine Learning (ML) models including Naive Bayes Classifier (NB), *K*-Nearest Neighbours Classifier (*K*NN), Random Forest Classifier (RF), Linear Support Vector Classifier (LSVC), Non-Linear Support Vector Classifier (NLSVC), and Regularized Logistic Regression Classifier (RLRC) using Python. First, we split up the dataset using a stratified (by class) test size of 1/3 (train_test_split function from scikit-learn Python module). Hyperparameters were optimized based on the model balanced accuracy (a more informative metric than simple accuracy, especially with imbalanced datasets) using a 10-fold Cross Validation (CV) approach (RepeatedStratifiedKFold, GridSearchCV, and balanced_accuracy functions from scikit-learn module). After tuning the hyperparameters, we compared the set of ML algorithms based on their best balance accuracy in a 10-times 10-fold CV framework. The best ML model was then trained with all the samples.

### Consensus Clustering of Tumor and Stroma Classifiers

The consensus clustering of both tumor and stroma classifiers consisted of two main steps: 1) network construction and 2) identification of consensus classes.

#### Tumor/Stroma Networks Construction

Once the samples were classified considering every classifier, we designed two binary matrices (with 1s if the sample belongs to the subtype *i* and 0s otherwise), each corresponding to the tumor and stroma classifications, whose dimensions were 514×19 and 514×4, respectively. Then, we constructed a network for each matrix using the igraph R package, where the edge weights were based on Cohen’s kappa index (κ^) estimated for each pairwise subtype comparison (subtypes *i* and *j*) using kappa2 function from irr R package. Subsequently, we applied a hypergeometric test using the phyper function from the stats R package to determine the probability of observing an equal or greater number of co-classifications between two different classes by chance. These p-values were corrected by multiple testing through False Discovery Rate (FDR) using the p.adjust function from the stats R package. Only edges with an FDR < 0.001 were considered.

#### Tumor/Stroma Consensus Classes Identification

We visually inspected the two generated clustering networks to identify the tumor/stroma consensus classes. This inspection allowed us to distinguish two well-defined consensus stroma clusters. However, given the difficulty of identifying consensus classes after inspecting the clustering network for the tumor classifiers, we applied a Markov Clustering Algorithm (MCL), especially indicated for clustering networks, using the MCL R package. It starts with an adjacent matrix representing the network, which is transformed into a Markov matrix where each element indicates the probability of moving from one node to another in a single step. It simulates random walks in the graph, refining them through an iterative process until convergence is reached. This iterative process depends on two parameters: expansion and inflation. The expansion involves raising the Markov matrix to a power (typically squaring it), which essentially simulates taking two steps in the graph instead of one, thereby considering paths of length two. Inflation involves raising each element of the Markov matrix to a specified power (the inflation factor *I*) and then re-normalizing each row to sum to one. This parameter highly influences the results, with a higher *I* value leading to more and smaller clusters. Thus, we evaluated the clustering provided by the MCL algorithm, considering values for *l* ranging from 1.5 to 5 with increments of 0.25. The robustness of the clustering for each *l* value was checked by performing 500 random resampling of 80% of the classes and computing the proportion of times two subtypes appeared in the same cluster, ending up with a total of 15 consensus matrices summarizing the results of the 500 resampling. We further clustered each consensus matrix using the hclust function from the stats R package with the Bray-Curtis (BC) distance computed with the vegdist function from the vegan R package and considering different numbers of clusters (*k* value ranging from 0 to 10). The optimal values for *k* and *I* parameters were those maximizing the average silhouette width (ASW), which measures how well each object lies within its cluster, calculated with the silhouette function from the cluster R package.

### Tumor and Stroma Consensus Classes Characterization

Once the consensus classes for tumor/stroma classifications were identified, we needed to characterize them by gaining insight into their molecular characteristics. To this end, we first identified the core samples that better define those consensus classes, i.e., those samples where all their assigned subtypes belonged to a specific consensus class. Then, we performed a differential expression analysis (DEA) with DESeq2 R package by using the batch-corrected gene counts of a random split of 80% of the core samples. To avoid overlooking genes that exhibit high expression in the less-represented class in each tumor/stroma consensus classification, we filtered out genes with fewer than five reads in 10%/40% of the samples. A final list of genes significantly differentially expressed (DE, adjusted p-value < 0.001) were sorted according to their absolute log2FC. A GSEA (over log2FC) using gseGO function against Biological Process (BP), Molecular Function (MF), and Cellular Component (CC) Gene Ontologies (GO), and gseKEGG function against Kyoto Encyclopedia of Genes and Genomes (KEGG) (clusterProfiler R package) was carried out.

### Tumor and Stroma Single Sample Consensus Classifier

The following pipeline was applied to develop the tumor/stroma single sample consensus classifiers. We kept the 80% of the core samples randomly selected previously and the significant genes resulting from the DEA to perform a feature selection (FS) process consisting of sequentially including the genes with the highest absolute log2FC among the overexpressed and the underexpressed genes into a RF model. The final number of selected features was determined in a 10-times 10-fold CV scenario using the RandomForestClassifier and RepeatedStratifiedKFold functions from the scikit-learn module after evaluating the model performance in terms of balanced accuracy. Once the FS process was finished, we evaluated the RF model performance on the remaining 20% of the core samples. To ensure that the balanced accuracy achieved with the final model is not overly dependent on the split made in core samples, we performed a 10-times 10-fold CV pipeline, including both the DEA and the FS steps. The estimated balanced accuracy of the final model was the average of the counterparts obtained for each time/fold.

Once the tumor/stroma single sample consensus classifiers were constructed, we classified the initial 514 PDAC samples and the tumors of the two external datasets: ICGC-au and PanGenEU-vs. In addition, a score representing the probability of belonging to the less-represented tumor/stroma consensus class was estimated for each PDAC sample.

### Relationships between Tumor & Stroma Classification

The relationship between tumor and stroma classifications was analyzed using a Chi-squared (χ^2^) test (chisq.test function from the stat R package). Additionally, the potential interdependencies between the continuous scores from the tumor (NonClassicalScore) and stroma (ActivatedECMScore) were examined using a linear model (lm function of the stats R package).

### Association between PDAC Consensus Classes and Overall Survival

We assessed the association of the PDAC consensus classes with overall survival using a multivariate Cox model adjusted for age, sex, and tumor stage.

### R package & Shiny app

To facilitate the use of our classification system for new PDAC samples, we developed an R package named *PDACMOC* (*PDACMolecularOmniClassifier*). It also enables users to obtain the subtypes defined by the classification systems included in developing this consensus classifier. It provides a plot of the final classifier balanced accuracy using ggplot2 R package. This package has been implemented in a user-friendly Shiny app to enhance accessibility and ease of use. For the development and implementation of the Shiny app, we utilized several R packages, including reticulate (to include Python code within R code), DT, shiny, shinydashboard, shinyjs, and shinythemes.

## RESULTS

### Study Population, Expression Data Retrieval & Preprocessing

**Table 1** shows the characteristics of the studies used to build and train the consensus classifier (except for the ICGC_v100, which did not have this data available). The median age was similar across all datasets. The sex distribution was nearly identical across datasets, with females representing around 45%–47%. However, significant differences in tumor stage were found (p.overall < 0.001), with stage 3 and 4 tumors being more prevalent in the PanGenEU dataset (12.7% and 10.8%, respectively) compared to the other datasets.

**Table 1.** Summary characteristics of the training cohort.

|  | TCGA | ICGC_v84 | PanGenEU | p.overall | p.TCGA vs<br>ICGC_v84 | p.TCGA vs<br>PanGenEU | p.ICGC_v84 vs<br>PanGenEU |
| --- | --- | --- | --- | --- | --- | --- | --- |
|  | *N=134* | *N=160* | *N=105* |  |  |  |  |
| Age | 66.0<br>[57.0;73.8] | 66.3<br>[57.5;73.8] | 67.1 [59.7;75.1] | 0.418 | 0.986 | 0.358 | 0.358 |
| Sex: |  |  |  | 0.984 | 1.000 | 1.000 | 1.000 |
| Female | 61 (45.5%) | 74 (46.2%) | 49 (46.7%) |  |  |  |  |
| Male | 73 (54.5%) | 86 (53.8%) | 56 (53.3%) |  |  |  |  |
| Stage: |  |  |  | <0.001 | 0.230 | 0.001 | <0.001 |
| 1 | 11 (8.21%) | 20 (12.5%) | 6 (5.88%) |  |  |  |  |
| 2 | 120 (89.6%) | 131 (81.9%) | 72 (70.6%) |  |  |  |  |
| 3 | 3 (2.24%) | 7 (4.38%) | 13 (12.7%) |  |  |  |  |
| 4 | 0 (0.00%) | 2 (1.25%) | 11 (10.8%) |  |  |  |  |

ICGC\_v84, International Cancer Genome Consortium Canada batch 1; TCGA; The Cancer Genome Atlas

The external validation studies (PanGenEU-vs and ICGC-au) showed comparable median age. Their sex distributions were also similar, with females representing 55.2% of the PanGenEU-vs and 50.0% of the ICGC-au datasets. However, they exhibited significant differences regarding tumor stage (p-value < 0.001) being ICGC-au enriched with stage 2 tumors (82.6% vs. 47.0% in PanGenEU-vs). Again, stage 3 and 4 tumors were more frequent in the PanGenEU-vs (39.4% and 12.1%) compared to ICGC-au (3.49% and 8.14%) (**Table 2**).

**Table 2.**
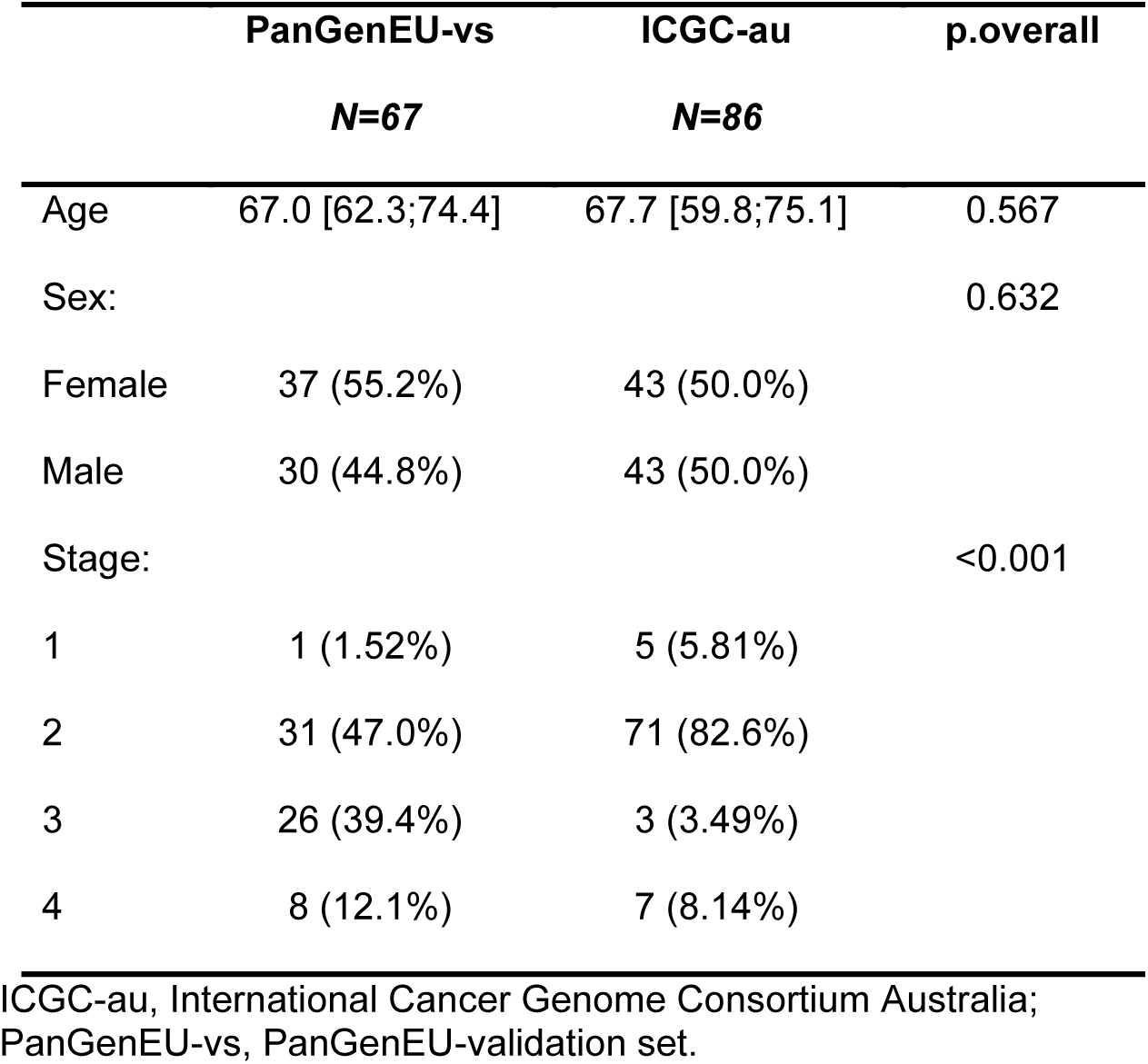
Summary characteristics table of the external cohort.

### Tumor and Stroma Single Sample Classifiers

We evaluated a set of ML models (i.e., NB, *K*NN, RF, LSVC, NLSVC, and RLRC), and the RLRC using Elastic-Net was the one with the largest balanced accuracy. Specifically, in the case of tumor samples, it was 97.4% for Collisson, 93.8% for Moffitt, 94.9% for Bailey, 97.9% for Puleo, and 98.4% for Chan-Seng-Yue. On the other hand, in the case of stroma samples, it was 98.9% for Moffitt and 91.1% for Maurer (**Supplementary Table 1**).

After classifying the individuals’ tumors in the training set, the distribution of the subtypes across different classifications is as follows (see **Figure 1** for further details): according to Collisson, 392 (76.3%) samples were classified as Classical, 22 (4.3%) as QM, and 100 (19.4%) as Exocrine-like. The Moffitt classification yielded 254 (49.4%) samples as Classical and 260 (50.6%) as Basal-like. For Bailey’s, there were 9 (1.8%) Squamous, 351 (68.3%) Progenitor, 87 (16.9%) Immunogenic, and 67 (13.0%) ADEX samples. Puleo’s classified 319 (62.1%) as Pure classical, 13 (2.5%) as Immune classical, 85 (16.5%) as Pure basal-like, 28 (5.5%) as Desmoplastic, and 69 (13.4%) as Stroma activated. Chan-Seng-Yue’s grouped 261 (50.8%) as Classical-A, 6 (1.2%) as Classical-B, 24 (4.6%) as Basal-A, 220 (42.8%) as Basal-B, and 3 (0.6%) as Hybrid. In the Moffitt stroma classification, 311 (60.5%) samples were classified as Normal and 203 (39.5%) as Activated, while the Maurer’s stroma classification identified 276 (53.7%) samples as ECM-rich and 238 (46.3%) as Immune-rich.

### Consensus Clustering of Tumor and Stroma Classifiers

The consensus clustering applied to the tumor classifiers detected two main groups: 1) Collisson Classical, Moffitt Classical, Bailey Progenitor, Puleo Pure Classical, Puleo Immune Classical, Chan-Seng-Yue Classical-A, and Chan-Seng-Yue Classical-B classes, and 2) Collisson QM, Collisson Exocrine-like, Moffitt Basal-like, Bailey Squamous, Bailey Immunogenic, Bailey ADEX, Puleo Pure basal-like, Puleo Desmoplastic, Puleo Stroma activated, Chan-Seng-Yue Basal-A, Chan-Seng-Yue Basal-B and Chan-Seng-Yue Hybrid classes (see **Figure 2a** for further details). We identified 159 core samples (i.e., samples where all their assigned subtypes belonged to a specific consensus class) of tumor consensus class 1 and 37 of tumor consensus class 2.

On the other hand, the consensus clustering applied to the stroma classifiers detected two groups (**Figure 2b**): 1) Moffitt Normal and Maurer Immune-rich subtypes, and 2) Moffitt Activated and Maurer ECM-rich subtypes. We detected 236 core samples of stroma consensus class 1 and 198 of stroma consensus class 2.

### Tumor and Stroma Consensus Classes Characterization

On the one hand, we identified 1,944 significant DE genes (1,081 overexpressed and 863 underexpressed) after comparing tumor consensus classes 1 and 2. The GSEA revealed that keratinocyte differentiation and cell adhesion genes were enriched in tumor consensus class 2 (see **Supplementary Figure 1** for further details). On the other hand, we identified 706 significant DE genes (516 overexpressed and 190 underexpressed) among stroma consensus classes 1 and 2. The GSEA revealed that genes related to the ECM were enriched in stroma consensus class 2 (see **Supplementary Figure 2** for further details).

Given the subtype composition of the tumor/stroma consensus classes and the GSEA results, we renamed the tumor consensus classes 1 and 2 as Consensus Classical and Consensus Non-classical, respectively, and the stroma consensus classes 1 and 2 as Consensus Normal-immune and Consensus Activated-ECM classes, respectively.

### Tumor and Stroma Single Sample Consensus Classifier

We evaluated different RF models to select the optimal number of significant DE genes to be included in the tumor single sample consensus classifier. Results showed that the RF, including 42 DE genes, yielded the best performance in terms of test-balanced accuracy (93.8%). This gene list includes 21 genes overexpressed in tumor samples classified as Consensus Classical (Classical signature) and 21 genes overexpressed in tumor samples classified as Consensus Non-classical (Non-classical signature). See **Table 3** for the complete signatures. We obtained a similar value when the DEA followed by the FS were carried out inside the 10-times 10-fold CV (averaged balance accuracy was 96.33% with a standard deviation of 7.88%). See **Table 4** for more details on the average balance accuracy obtained for the tumor SSCs.

**Table 3.** Gene signatures characterizing the Consensus Classical/Non-classical classes according to the tumor consensus classifier.

| **Classical** | **Non-classical** |
| --- | --- |
| **signature** | **signature** |
| *MUC2* | *FAT2* |
| *SPINK4* | *KRT5* |
| *TM4SF20* | *CPA1* |
| *KRT20* | *CTRB2* |
| *B3GNT6* | *CPB1* |
| *REG4* | *CELA3A* |
| *BTNL3* | *CTRB1* |
| *HMGCS2* | *CPA2* |
| *GPA33* | *REG1B* |
| *ADGRG7* | *CTRC* |
| *ITLN1* | *AMY2A* |
| *GUCY2C* | *CEL* |
| *MUC17* | *KRT14* |
| *SLC13A2* | *DNER* |
| *PRAP1* | *TNNT3* |
| *FEZF1* | *RHCG* |
| *BRNL8* | *H19* |
| *APOBEC1* | *TENM2* |
| *TFF1* | *KRT13* |
| *PHGR1* | *KRT6A* |
| *MYO7B* | *IGKV1-12* |

**Table 4.** Mean balanced accuracy of the final tumor single sample classifiers.

| Classifier | Balanced accuracy (%) |
| --- | --- |
| Collisson (RLRC) | 97.44 |
| Moffitt (RLRC) | 93.78 |
| Bailey (RLRC) | 94.95 |
| Puleo (RLRC) | 97.95 |
| Chan-Seng-Yue (RLRC) | 98.42 |
| **PDAConsensus (RF)** | **96.33** |

RF, Random Forest Classifier; RLRC, Regularized Logistic Regression Classifier.

Results of the evaluation of different RF models to select the optimal number of significant DE genes to be included in the stroma single sample consensus classifier showed that the RF, including 60 DE genes, yielded the best performance in terms of the test balanced accuracy (100%). This gene list includes 30 genes overexpressed in stroma samples classified as Consensus Normal-immune (Normal-immune signature) and 30 genes overexpressed in stroma samples classified as Consensus Activated-ECM (Activated-ECM signature). See **Table 5** for the complete signatures. A slightly lower balanced accuracy was obtained when both the DEA and the FS were performed inside the 10-times 10-fold CV (mean balanced accuracy of 98.92% with a standard deviation of 1.71%. See **Table 6** for more details on the average balance accuracy obtained for the stroma SSCs.

**Table 5.** Gene signatures characterizing the Consensus Normal-immune and the Activated-ECM classes according to the stroma consensus classifier.

| Normal-immune | Activated-ECM |
| --- | --- |
| signature | signature |
| *MUC2* | *EPYC* |
| *SPINK4* | *KRT14* |
| *VTN* | *LRRC15* |
| *SLC6A19* | *OMD* |
| *APOB* | *MFAP5* |
| *PGC* | *SFRP2* |
| *ITLN1* | *ADAM12* |
| *SLC28A2* | *IGKV1-12* |
| *GP2* | *COL11A1* |
| *ALB* | *FAM180A* |
| *PSAPL1* | *FNDC1* |
| *PDIA2* | *GALNT15* |
| *REG4* | *POSTN* |
| *REG1B* | *COL10A1* |
| *A4GNT* | *FIBIN* |
| *C2CD4B* | *COL1A1* |
| *SLC13A2* | *WT1* |
| *AKR1C4* | *COL1A2* |
| *SLC19A3* | *CMKLR2* |
| *TFF2* | *TIMP3* |
| *MT1H* | *XIRP1* |
| *REG3A* | *COL3A1* |
| *PRAP1* | *MEDAG* |
| *MAP3K20-AS1* | *MAGEL2* |
| *SLC39A5* | *FN1* |
| *TM4SF20* | *COL6A3* |
| *MT1G* | *IGFL2* |
| *HPN* | *ST6GAL2* |
| *SH2D6* | *HSD17B6* |
| *POU2AF2* | *GREM1* |

**Table 6.** Mean balanced accuracy of the final stroma singe sample classifiers.

| Classifier | Balanced accuracy (%) |
| --- | --- |
| Moffitt (RLRC) | 98.86 |
| Maurer (RLRC) | 91.10 |
| **PDAConsensus (RF)** | **98.92** |

RF, Random Forest Classifier; RLRC, Regularized Logistic Regression Classifier.

Overall, our single sample consensus classifiers utilize 903 distinct genes for tumor and 532 distinct genes for stroma grouping, amounting to a total of 1,330 distinct genes (105 shared between them). Among the 42 genes identified for the Tumor Consensus classifier, 35 were present in other classifiers, with *B3GNT6*, *FEZF1, PRAP1*, *FAM177B*, *SLC13A2*, *DNER*, *H19*, and *TENM2* being the exceptions (**Supplementary Table 2**).

We explored the single-cell expression of those seven genes using the web tool (https://sunshine.bioinformatics.cnio.es/groups/carcinogenesis/Human_PDAC/) and corroborated that they follow a similar pattern, confirming their validity and correctness (**Supplementary Figure 3**). Of note, *GATA6*, an epithelial gene known as a classical marker used in immunohistochemistry studies, was not selected among the top DE genes; therefore, it was not selected by the RF in the FS step. However, *GATA6* was indeed overexpressed in Consensus Classical compared to Consensus Non-classical samples (log2FC = −1.8974, adjusted p-value < 0.001) and ranked downstream.

In contrast, among the 60 genes identified for the stroma consensus classifier, only 16 are present in other classifiers (*ADAM12*, *COL10A1*, *COL11A1*, *COL1A1*, *COL1A2*, *COL3A1*, *COL6A3*, *EPYC*, *FN1*, *FNDC1*, *GREM1*, *LRRC15*, *MFAP5*, *POSTN*, *SFRP2*, and *TIMP3*).

**Figures 3a** and **3b** present parallel sets illustrating how samples classified according to the tumor/stroma consensus classifiers are categorized by previously published classifiers. These visualizations provide insights into the alignment and variation in subtype classifications across different frameworks.

### Relationship between Tumor & Stroma Consensus Classifications

The result of the χ2-test suggested an association between the tumor and stroma consensus classifications (p-value = 0.05). Specifically, if a tumor is classified as Consensus Classical, the estimated probability of the stroma being Consensus Normal-immune is 59.6%, whereas the likelihood of Consensus Activated-ECM stroma in the presence of a Consensus Non-classical tumor is 50.4%. Conversely, the probability of a Consensus Classical tumor given Consensus Normal-immune stroma is 75.7%, and the likelihood of a Consensus Non-classical tumor in the case of Consensus Activated-ECM stroma is 32.4%. Figure 4 illustrates a positive linear relationship between NonClassicalScore and ActivatedECMScore, as captured by the regression equation y=0.21x+0.38 (p-value < 0.001), suggesting that an increase in NonClassicalScore is associated with an increase in ActivatedECMScore. The marginal density plots emphasize the score distributions: the NonClassicalScore predominantly shows lower values, which is consistent with a higher prevalence of samples classified as Consensus Classical. In contrast, the ActivatedECMScore distribution is bimodal, mirroring the equilibrium observed among the stroma subtypes.

**Figure 4.**
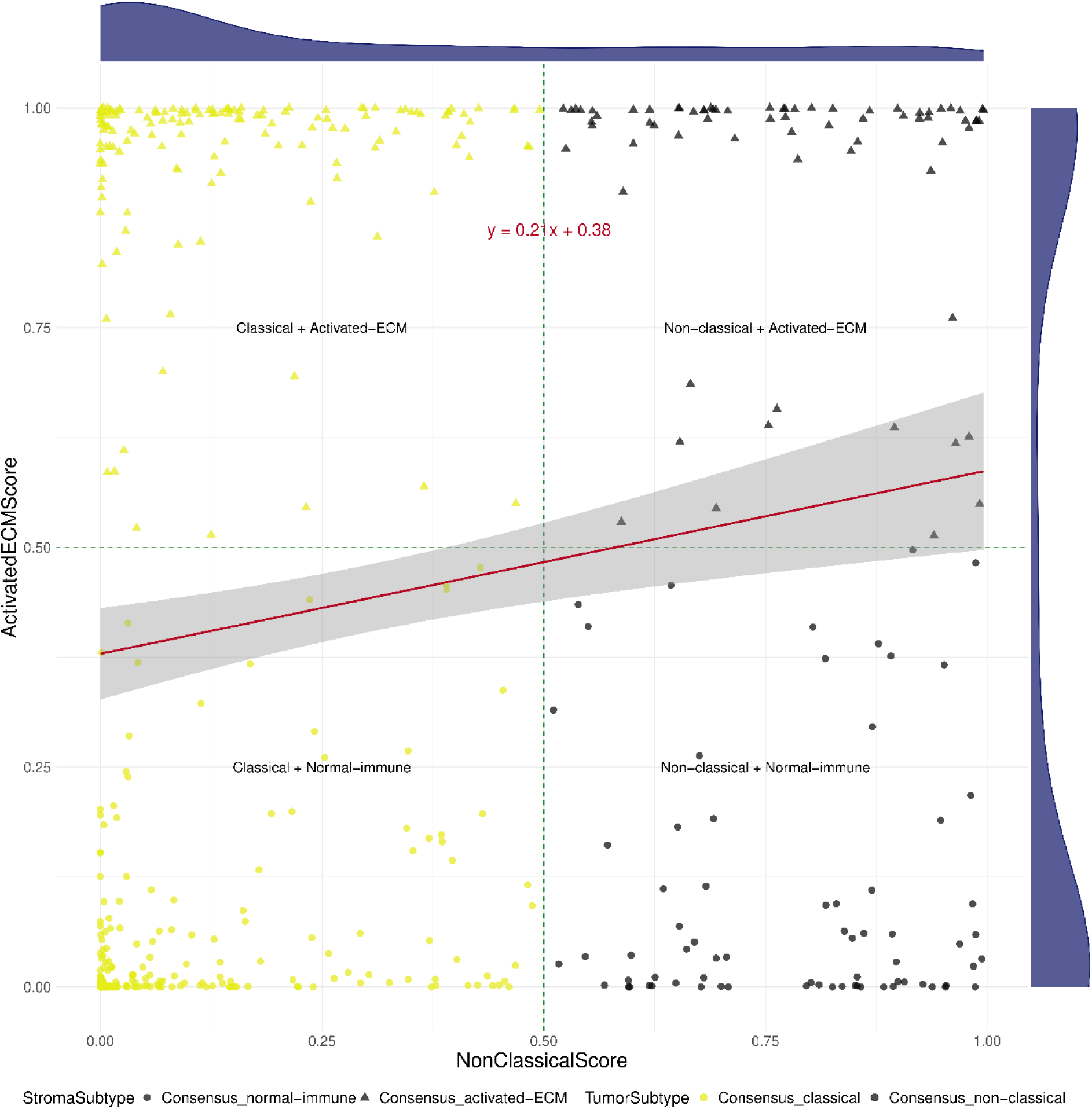
Correlation between NonClassicalScore and ActivatedECMScore.

### Association between PDAC Consensus Classes and Overall Survival

The association between the SSCs of the tumor/stroma and overall survival was tested in the validation datasets (Figure 5) and the training datasets (**Supplementary Figure 4**). We found a significant association between the tumor consensus classes and overall survival in TCGA and ICGC-au studies. Results show that patients with tumors classified as Consensus Non-classical had a higher risk of death than those with Consensus Classical tumors. However, the associations were not statistically significant in ICGC_v84 (ICGC Canada) and PanGenEU. Results from the other classification systems were heterogeneous.

**Figure 5.**
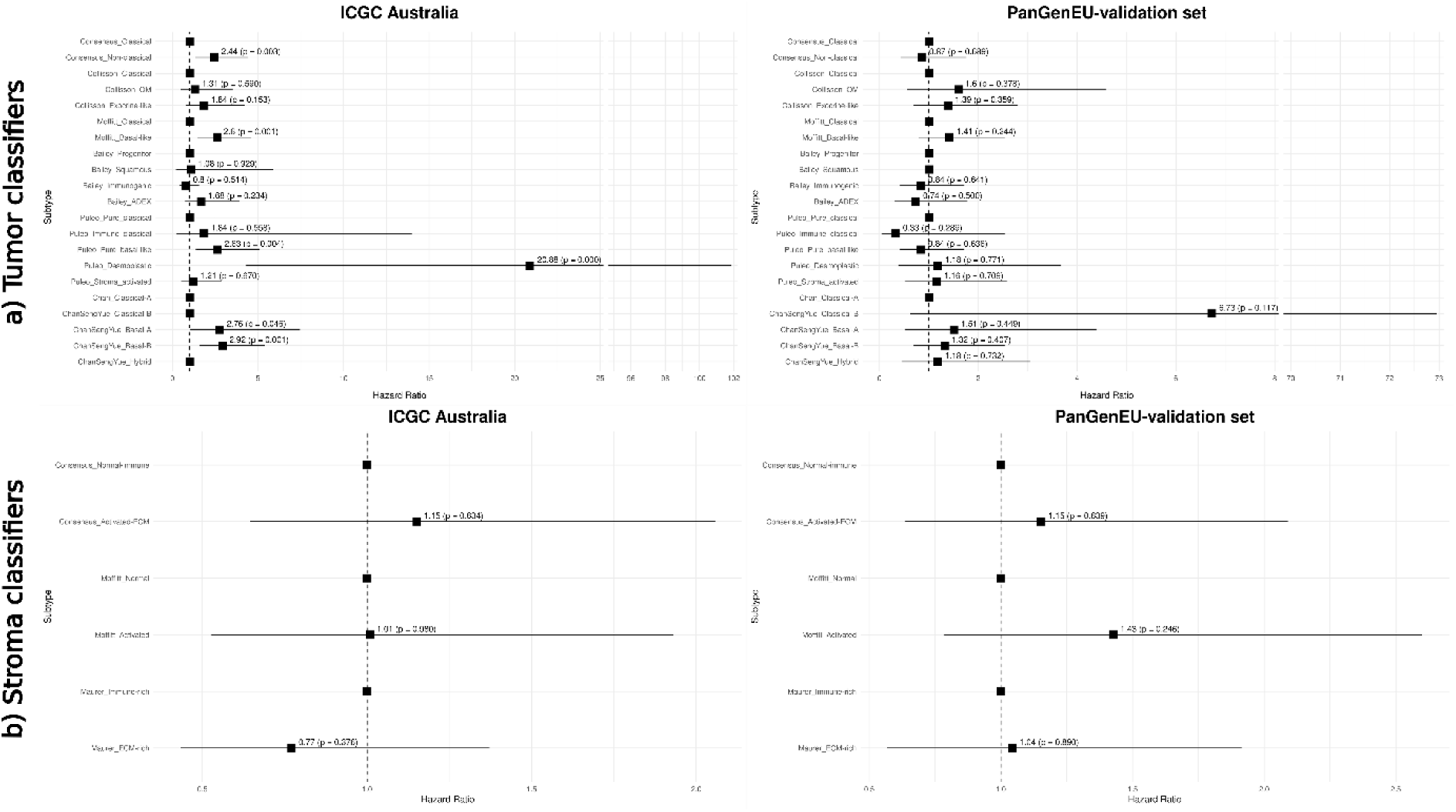
Cox Proportional Hazards models of tumor and stroma classifiers in the validation datasets. ADEX (aberrantly differentiated endocrine-exocrine), ECM (extracellular matrix), ICGC (International Cancer Genome Consortium), QM (quasi-mesenchymal.

Results of the association between stroma consensus classes and overall survival showed that patients with Consensus Activated-ECM tumors survived less than those whose tumors were classified as Consensus Normal-immune in ICGC_v84 (ICGC Canada. The associations were not statistically significant in TCGA, ICGC-au, and PanGenEU datasets. Again, results from the other classification systems were heterogeneous.

## DISCUSSION

By employing cutting-edge ML methods applied to a large dataset comprising 514 PDAC tumors, we developed a consensus classifier (*PDAConsensus*) that integrates information from several previously established molecular classifiers, resulting in a binary classification for both tumor and stroma. The consensus classifier categorized both the tumor and stroma compartments into two robust classes each (Consensus Classical and Consensus Non-classical, and Consensus Normal-immune and Consensus Activated-ECM, respectively), providing a more nuanced understanding of the heterogeneous nature of PDAC. We also constructed a NonClassicalScore and an ActivatedECMScore to characterize the tumor and stroma compartments better. This dual-component approach is pivotal given the growing recognition of the stroma’s role in cancer biology, particularly in PDAC (5–9). Additionally, we have demonstrated the association of tumor and stroma classes provided by the *PDAConsensus* classifier with overall survival in TCGA and ICGC-au despite the association lacking significance in the remaining datasets. These discrepancies could be mainly attributed to inherent differences between the datasets, such as the tumor stage and PDAC patient management in distinct clinical settings. Nevertheless, our consensus classifier allows us to classify the PDAC samples based on their tumor/stroma molecular characteristics and stratify the PDAC patients based on their overall survival at least as effectively as other classifiers.

The stroma plays a crucial role in PDAC and must be integrated into classification systems like our consensus classifier. Interestingly, we observed an association between the tumor and stroma consensus classifications and a linear relationship between the NonClassicalScore and the ActivatedECMScore. These findings align with Oh et al., who used single-cell RNA sequencing data to correlate tumor subtypes (classical and basal-like) with distinct TME activities (23).

Our tumor gene signatures demonstrate strong validation based on their overlap with established classification systems, as detailed in **Supplementary Table 2**. This alignment emphasizes our FS process’s robustness and the identified genes’ relevance. We further validated the expression patterns of the unique tumor genes in our classifier through single-cell analysis, thereby confirming their biological plausibility. Finally, while *GATA6* did not rank among the top genes selected during the classification process, we confirmed its overexpression in Consensus Classical samples compared to Consensus Non-classical samples.

This study employed a comprehensive and state-of-the-art methodological framework for developing and validating a consensus molecular classifier for PDAC. Using mRNA expression data from multiple international datasets (TCGA, ICGC, and PanGenEU), robust preprocessing steps, including batch correction, low-expression filtering, and VST, ensured data harmonization. Advanced ML techniques were applied to train and validate tumor and stroma SSCs. Innovative tools like the MCL were employed, and stratified CV pipelines were implemented to enhance classifier accuracy and robustness. To promote accessibility and usability, the *PDACMOC* R package and a user-friendly Shiny app were developed, allowing seamless application of these classifiers to new PDAC samples and fostering broader adoption in translational research.

While the present study’s findings are promising, certain limitations merit consideration. One initial and most significant step involved classifying each sample according to each available classifier. However, since most of these classifiers were developed using clustering methods, classifying a single new sample is not feasible. Consequently, we had to define a “ground truth” to create an SSC for each classification system. This step is crucial, as all subsequent work depends on it. For these tasks, we opted for hierarchical clustering combined with GSEA, except for Puleo’s classifier, where we applied correlation to centroids as suggested by the authors. Another vital aspect is VM. Although there are numerous methods for this, most are based on clustering and are unsuitable for our approach. Therefore, we ultimately chose the ADVOCATE R package.

Extensive work and data will validate this classifier in additional clinical cohorts by exploring its predictive power for tumor growth and treatment response by integrating other data layers. Comparisons with classifiers based on metabolic (24), morphological (25), splicing (26), and endoplasmic reticulum stress/lipid metabolism/liver metastasis (27) subtypes will help identify similarities and differences among subtypes. Expanding beyond bulk RNA data to include single-cell RNA sequencing, single-nucleus RNA sequencing, and spatial transcriptomics could significantly enhance our understanding of PDAC heterogeneity and tumor dynamics. Integrating advanced methods such as deep learning and neural networks to differentiate tumor from stroma components may further improve model accuracy. Additionally, identifying consensus subtypes using minimally invasive techniques like liquid biopsy would greatly benefit patients, as acknowledged by Metzenmacher et al. (28). Enhancing the classifier with new data and incorporating multi-omics strategies will guarantee its ongoing relevance and usefulness in clinical practice.

The consensus classifier offers a comprehensive and integrated approach to PDAC classification, tackling the challenge of using multiple existing systems. Its clinical implications may be significant, allowing customized treatment plans, enhancing prognostic evaluations, and directing the creation of targeted therapeutic agents. By ensuring more homogenous patient groups in clinical trials, the classifier enhances the reliability of results and promotes precision medicine, ultimately improving patient outcomes.

In summary, the development of *PDAConsensus* represents a significant advancement in PDAC research. Its capacity to incorporate both tumor and stroma components could transform our understanding and management of PDAC. By making this technology accessible through the package *PDACMOC* R and a Shiny app, this study paves the way for its broader application in understanding and patient management of PDAC.

## DATA AVAILABILITY

The *PDACMOC* R package is available at GitHub (https://github.com/pavillos/PDACMOC) and the Shiny app can be accessed at https://pdacmoc.cnio.es/. Supplementary Data are available at bioRxiv online.

## AUTHOR CONTRIBUTIONS

*Pablo Villoslada-Blanco*: Conceptualization, Formal analysis, Methodology, Validation, Writing—original draft, review & editing. *Lola Alonso*: Data acquisition and custody, Methodology, Validation, Writing—review. *Sergio Sabroso-Lasa*: Validation, Writing— review. *Miguel Maquedano*: Formal analysis, Writing—review. *Lidia Estudillo*: Sample collection, Extraction protocol design, and Sample processing. *Francisco X Real*: Conceptualization, Writing—review. *Evangelina López de Maturana*: Conceptualization, Validation, Writing—review. *Núria Malats*: Conceptualization, Funding, Methodology, Validation, Writing—review.

## ACKNOWLEDGEMENTS

The authors thank the patients, coordinators, field and administrative workers, and technicians of the European Study into Digestive Illnesses and Genetics (PanGenEU) study.

## ABBREVIATIONS

ADEX: aberrantly differentiated endocrine-exocrine
ASW: average silhouette width
BC: Bray-Curtis
BP: Biological Process
CAFs: cancer associated fibroblasts
CC: Cellular Component
CV: Cross Validation
DEA: differential expression analysis
ECM: extracellular matrix
FDR: False Discovery Rate
FS: feature selection
GO: Gene Ontologies
GSEA: gene set enrichment analysis
ICGC: International Cancer Genome Consortium
ICGC-au: International Cancer Genome Consortium Australia
ICGC_v100: International Cancer Genome Consortium Canada batch 1
ICGC_v84: International Cancer Genome Consortium Canada batch 2
KEGG: Kyoto Encyclopedia of Genes and Genomes
*K*NN: *K*-Nearest Neighbours Classifier
LSVC: Linear Support Vector Classifier
MCL: Markov Clustering Algorithm
MF: Molecular Function
ML: Machine Learning
NB: Naive Bayes Classifier
NLSVC: Non-Linear Support Vector Classifier
PanGenEU-vs: PanGenEU validation set
PDAC: Pancreatic Ductal Adenocarcinoma
PDACMOC: PDACMolecularOmniClassifier
QM: quasi-mesenchymal
RF: Random Forest Classifier
RLRC: Regularized Logistic Regression Classifier
SSC: single sample classifier
TCGA: The Cancer Genome Atlas
TME: tumor microenvironment
VM: virtual microdissection
VST: Variance-Stabilizing Transformation

## SUPPLEMENTARY TABLES AND FIGURES

**Supplementary Table 1.** Balanced accuracy of the different Machine Learning algorithms.

**Supplementary Table 2**. Presence of the genes belonging to the Classical signature and the Non-classical signature in the other classification systems.

**Supplementary Figure 1.** Gene Set Enrichment Analysis of tumor consensus classes. GeneRatio measures the proportion of genes in a particular gene set that are found within the ranked lists of genes, while enrichment distribution quantifies the degree to which a gene set is overrepresented at the top or bottom of a ranked list of genes. GO, Gene Ontology; KEGG, Kyoto Encyclopedia of Genes and Genomes.

**Supplementary Figure 2**. Gene Set Enrichment Analysis of stroma consensus classes. GeneRatio is a measure of the proportion of genes in a particular gene set that are found within the ranked lists of genes, while enrichment distribution quantifies the degree to which a gene set is overrepresented at the top or bottom of a ranked list of genes. GO, Gene Ontology; KEGG, Kyoto Encyclopedia of Genes and Genomes.

**Supplementary Figure 3**. Expression of the genes belonging to the Classical and Non-classical signatures in a single-cell dataset. At the top, as a reference, the different clusters and the expression of GATA6 are shown. Data for H19 was not available.

**Supplementary Figure 4.** Cox Proportional Hazards models of tumor and stroma classifiers in the training cohort. All the models are adjusted by Age, Sex and Stage, and Hazar Ratio and p-value are presented. ADEX (aberrantly differentiated endocrine-exocrine), ECM (extracellular matrix), ICGC (International Cancer Genome Consortium), QM (quasi-mesenchymal), TCGA (The Cancer Genome Atlas).

## REFERENCES

1. Siegel RL, Miller KD, Wagle NS, Jemal A. Cancer statistics, 2023. CA Cancer J Clin [Internet]. 2023 Jan [cited 2023 Nov 14];73(1):17–48. Available from: https://pubmed.ncbi.nlm.nih.gov/36633525/

2. Mizrahi JD, Surana R, Valle JW, Shroff RT. Pancreatic cancer. Lancet (London, England) [Internet]. 2020 Jun 27 [cited 2023 Dec 20];395(10242):2008–20. Available from: https://pubmed.ncbi.nlm.nih.gov/32593337/

3. Bailey P, Chang DK, Nones K, Johns AL, Patch AM, Gingras MC, et al. Genomic analyses identify molecular subtypes of pancreatic cancer. Nature [Internet]. 2016 Mar 3 [cited 2023 Nov 14];531(7592):47–52. Available from: https://pubmed.ncbi.nlm.nih.gov/26909576/

4. Hoadley KA, Yau C, Hinoue T, Wolf DM, Lazar AJ, Drill E, et al. Cell-of-Origin Patterns Dominate the Molecular Classification of 10,000 Tumors from 33 Types of Cancer. Cell [Internet]. 2018 Apr 5 [cited 2023 Nov 14];173(2):291–304.e6. Available from: https://pubmed.ncbi.nlm.nih.gov/29625048/

5. Erkan M, Hausmann S, Michalski CW, Fingerle AA, Dobritz M, Kleeff J, et al. The role of stroma in pancreatic cancer: diagnostic and therapeutic implications. Nat Rev Gastroenterol Hepatol [Internet]. 2012 Aug [cited 2023 Oct 14];9(8):454–67. Available from: https://pubmed.ncbi.nlm.nih.gov/22710569/

6. Whatcott CJ, Diep CH, Jiang P, Watanabe A, Lobello J, Sima C, et al. Desmoplasia in Primary Tumors and Metastatic Lesions of Pancreatic Cancer. Clin Cancer Res [Internet]. 2015 Aug 1 [cited 2023 Oct 14];21(15):3561–8. Available from: https://pubmed.ncbi.nlm.nih.gov/25695692/

7. Elyada E, Bolisetty M, Laise P, Flynn WF, Courtois ET, Burkhart RA, et al. Cross-Species Single-Cell Analysis of Pancreatic Ductal Adenocarcinoma Reveals Antigen-Presenting Cancer-Associated Fibroblasts. Cancer Discov [Internet]. 2019 Aug 1 [cited 2023 Oct 14];9(8):1102–23. Available from: https://pubmed.ncbi.nlm.nih.gov/31197017/

8. Feig C, Gopinathan A, Neesse A, Chan DS, Cook N, Tuveson DA. The pancreas cancer microenvironment. Clin Cancer Res [Internet]. 2012 Aug 15 [cited 2023 Oct 14];18(16):4266–76. Available from: https://pubmed.ncbi.nlm.nih.gov/22896693/

9. Biffi G, Oni TE, Spielman B, Hao Y, Elyada E, Park Y, et al. IL1-Induced JAK/STAT Signaling Is Antagonized by TGFβ to Shape CAF Heterogeneity in Pancreatic Ductal Adenocarcinoma. Cancer Discov [Internet]. 2019 [cited 2023 Oct 14];9(2):282–301. Available from: https://pubmed.ncbi.nlm.nih.gov/30366930/

10. Collisson EA, Sadanandam A, Olson P, Gibb WJ, Truitt M, Gu S, et al. Subtypes of pancreatic ductal adenocarcinoma and their differing responses to therapy. Nat Med [Internet]. 2011;17(4):500–3. Available from: 10.1038/nm.2344

11. Moffitt RA, Marayati R, Flate EL, Volmar KE, Loeza SGH, Hoadley KA, et al. Virtual microdissection identifies distinct tumor- and stroma-specific subtypes of pancreatic ductal adenocarcinoma. Nat Genet [Internet]. 2015;47(10):1168–78. Available from: 10.1038/ng.3398

12. Puleo F, Nicolle R, Blum Y, Cros J, Marisa L, Demetter P, et al. Stratification of Pancreatic Ductal Adenocarcinomas Based on Tumor and Microenvironment Features. Gastroenterology [Internet]. 2018 Dec 1;155(6):1999–2013.e3. Available from: 10.1053/j.gastro.2018.08.033

13. Chan-Seng-Yue M, Kim JC, Wilson GW, Ng K, Figueroa EF, O’Kane GM, et al. Transcription phenotypes of pancreatic cancer are driven by genomic events during tumor evolution. Nat Genet [Internet]. 2020;52(2):231–40. Available from: 10.1038/s41588-019-0566-9

14. Maurer C, Holmstrom SR, He J, Laise P, Su T, Ahmed A, et al. Experimental microdissection enables functional harmonisation of pancreatic cancer subtypes. Gut [Internet]. 2019 Jun 1;68(6):1034. Available from: http://gut.bmj.com/content/68/6/1034.abstract

15. Collisson EA, Bailey P, Chang DK, Biankin A V. Molecular subtypes of pancreatic cancer. Vol. 16, Nature Reviews Gastroenterology and Hepatology. Nature Publishing Group; 2019. p. 207–20.

16. Guinney J, Dienstmann R, Wang X, De Reyniès A, Schlicker A, Soneson C, et al. The consensus molecular subtypes of colorectal cancer. Nat Med. 2015 Nov 1;21(11):1350–6.

17. Kamoun A, de Reyniès A, Allory Y, Sjödahl G, Robertson AG, Seiler R, et al. A Consensus Molecular Classification of Muscle-invasive Bladder Cancer. Eur Urol [Internet]. 2020;77(4):420–33. Available from: https://www.sciencedirect.com/science/article/pii/S0302283819306955

18. López de Maturana E, Rodríguez JA, Alonso L, Lao O, Molina-Montes E, Martín- Antoniano IA, et al. A multilayered post-GWAS assessment on genetic susceptibility to pancreatic cancer. Genome Med [Internet]. 2021;13(1):15. Available from: 10.1186/s13073-020-00816-4

19. Molina-Montes E, Coscia C, Gómez-Rubio P, Fernández A, Boenink R, Rava M, et al. Deciphering the complex interplay between pancreatic cancer, diabetes mellitus subtypes and obesity/BMI through causal inference and mediation analyses. Gut. 2021;70(2):319–29.

20. Zhang Y, Parmigiani G, Johnson WE. ComBat-seq: batch effect adjustment for RNA-seq count data. NAR Genomics Bioinforma [Internet]. 2020 Sep 1 [cited 2024 Jul 7];2(3). Available from: 10.1093/nargab/lqaa078

21. Love MI, Huber W, Anders S. Moderated estimation of fold change and dispersion for RNA-seq data with DESeq2. Genome Biol [Internet]. 2014 Dec 5 [cited 2024 Jul 8];15(12):1–21. Available from: https://genomebiology.biomedcentral.com/articles/10.1186/s13059-014-0550-8

22. He J, Maurer HC, Holmstrom SR, Su T, Ahmed A, Hibshoosh H, et al. Transcriptional deconvolution reveals consistent functional subtypes of pancreatic cancer epithelium and stroma. bioRxiv [Internet]. 2018 Mar 27 [cited 2023 Nov 16];288779. Available from: https://www.biorxiv.org/content/10.1101/288779v3

23. Oh K, Yoo YJ, Torre-Healy LA, Rao M, Fassler D, Wang P, et al. Coordinated single-cell tumor microenvironment dynamics reinforce pancreatic cancer subtype. Nat Commun [Internet]. 2023 Aug 26;14(1):5226. Available from: http://www.ncbi.nlm.nih.gov/pubmed/37633924

24. Pervin J, Asad M, Cao S, Jang GH, Feizi N, Haibe-Kains B, et al. Clinically impactful metabolic subtypes of pancreatic ductal adenocarcinoma (PDAC). Front Genet [Internet]. 2023 [cited 2023 Dec 4];14. Available from: /pmc/articles/PMC10643182/

25. Kalimuthu SN, Wilson GW, Grant RC, Seto M, O’Kane G, Vajpeyi R, et al. Morphological classification of pancreatic ductal adenocarcinoma that predicts molecular subtypes and correlates with clinical outcome. Gut [Internet]. 2020 Feb 1;69(2):317. Available from: http://gut.bmj.com/content/69/2/317.abstract

26. Ruta V, Naro C, Pieraccioli M, Leccese A, Archibugi L, Cesari E, et al. An alternative splicing signature defines the basal-like phenotype and predicts worse clinical outcome in pancreatic cancer. Cell reports Med [Internet]. 2024 Feb [cited 2024 Mar 18];5(2):101411. Available from: https://pubmed.ncbi.nlm.nih.gov/38325381/

27. Liu X, Ren B, Fang Y, Ren J, Wang X, Gu M, et al. Comprehensive analysis of bulk and single-cell transcriptomic data reveals a novel signature associated with endoplasmic reticulum stress, lipid metabolism, and liver metastasis in pancreatic cancer. J Transl Med [Internet]. 2024 Dec 1 [cited 2024 May 7];22(1). Available from: https://pubmed.ncbi.nlm.nih.gov/38685045/

28. Metzenmacher M, Zaun G, Trajkovic-Arsic M, Cheung P, Reissig TM, Schürmann H, et al. Minimally invasive determination of pancreatic ductal adenocarcinoma (PDAC) subtype by means of circulating cell-free RNA. Mol Oncol [Internet]. 2024 Oct 31; Available from: http://www.ncbi.nlm.nih.gov/pubmed/39478658

