## Supplementary material for "Development of a Consensus Molecular Classifier for Pancreatic Ductal Adenocarcinoma"

Pablo Villoslada-Blanco *et al*.


| | |
|---|---|
| **Supplementary Table 1** | Page 2 |
| **Supplementary Table 2** | Page 3 |
| **Supplementary Figure 1** | Page 5 |
| **Supplementary Figure 2** | Page 6 |
| **Supplementary Figure 3** | Page 7 |
| **Supplementary Figure 4** | Page 8 |

**Supplementary Table 1**. Balanced accuracy of the different Machine Learning algorithms.

| | | NB | *K*NN | RF | LSVC | NLSVC | RLRC |
|---|---|---|---|---|---|---|---|
| **TUMOR CLASSIFIERS** | Collisson | 0.9585 | 0.8442 | 0.8470 | 0.9234 | 0.9122 | 0.9744 |
| | Moffitt | 0.8429 | 0.8999 | 0.9046 | 0.8934 | 0.8938 | 0.9378 |
| | Bailey | 0.8039 | 0.7795 | 0.6186 | 0.7490 | 0.7920 | 0.9495 |
| | Puleo | 0.8313 | 0.6870 | 0.6098 | 0.8538 | 0.8568 | 0.9795 |
| | Chan-Seng-Yue | 0.8514 | 0.8221 | 0.7673 | 0.8889 | 0.8891 | 0.9842 |
| **STROMA CLASSIFIERS** | Moffitt | 0.9542 | 0.9359 | 0.9680 | 0.9679 | 0.9658 | 0.9886 |
| | Maurer | 0.8827 | 0.9140 | 0.9068 | 0.8973 | 0.9007 | 0.9110 |

*K*NN (*K*-Nearest Neighbour Classifier), LSVC (Linear Support Vector Classifier), NB (Naive Bayes Classifier), NLSVC (Non-Lineal Support Vector Classifier), RF (Random Forest Classifier), RLRC (Regularized Logistic Regression Classifier).

**Supplementary Table 2**. Genes belonging to the Classical and Non-Classical signatures present in other classification systems.

| | | Collisson_Classical | Moffitt_Classical | Bailey_Progenitor | Chan-Seng-Yue_Classical-A | Chan-Seng-Yue_Classical-B | Collisson_QM | Moffitt_Basal-like | Bailey_Squamous | Chan-Seng-Yue_Basal-A | Chan-Seng-Yue_Basal-B | Collisson_Exocrine-like | Bailey_ADEX | Bailey_Immunogenic | Puleo | Consensus_classical | Consensus_non-classical |
|---|---|---|---|---|---|---|---|---|---|---|---|---|---|---|---|---|---|
| | GATA6 | 0 | 0 | 0 | 1 | 0 | 0 | 0 | 0 | 0 | 0 | 0 | 0 | 0 | 0 | 0 | 0 |
| **Typical classical genes** | ADGRG7 | 0 | 0 | 0 | 0 | 1 | 0 | 0 | 0 | 0 | 0 | 0 | 0 | 0 | 0 | 1 | 0 |
| | APOBEC1 | 0 | 0 | 0 | 1 | 0 | 0 | 0 | 0 | 0 | 0 | 0 | 0 | 0 | 0 | 1 | 0 |
| | BTNL3 | 0 | 0 | 0 | 0 | 1 | 0 | 0 | 0 | 0 | 0 | 0 | 0 | 0 | 0 | 1 | 0 |
| | BTNL8 | 0 | 1 | 1 | 0 | 1 | 0 | 0 | 0 | 0 | 0 | 0 | 0 | 0 | 0 | 1 | 0 |
| | GPA33 | 0 | 0 | 0 | 0 | 1 | 0 | 0 | 0 | 0 | 0 | 0 | 0 | 0 | 0 | 1 | 0 |
| | GUCY2C | 0 | 0 | 0 | 0 | 1 | 0 | 0 | 0 | 0 | 0 | 0 | 0 | 0 | 0 | 1 | 0 |
| | HMGCS2 | 0 | 0 | 0 | 0 | 1 | 0 | 0 | 0 | 0 | 0 | 0 | 0 | 0 | 1 | 1 | 0 |
| | ITLN1 | 0 | 0 | 0 | 0 | 1 | 0 | 0 | 0 | 0 | 0 | 0 | 0 | 1 | 0 | 1 | 0 |
| | KRT20 | 0 | 1 | 0 | 0 | 1 | 0 | 0 | 0 | 0 | 0 | 0 | 0 | 0 | 0 | 1 | 0 |
| | MUC2 | 0 | 0 | 0 | 0 | 1 | 0 | 0 | 0 | 0 | 0 | 0 | 0 | 0 | 0 | 1 | 0 |
| | MUC17 | 0 | 0 | 1 | 1 | 0 | 0 | 0 | 0 | 0 | 0 | 0 | 0 | 0 | 1 | 1 | 0 |
| | MYO7B | 0 | 0 | 1 | 0 | 1 | 0 | 0 | 0 | 0 | 0 | 0 | 0 | 0 | 0 | 1 | 0 |
| | PHGR1 | 0 | 0 | 1 | 0 | 1 | 0 | 0 | 0 | 0 | 0 | 0 | 0 | 0 | 0 | 1 | 0 |
| | REG4 | 0 | 1 | 1 | 0 | 1 | 0 | 0 | 0 | 0 | 0 | 0 | 0 | 0 | 0 | 1 | 0 |
| | SPINK4 | 0 | 1 | 0 | 0 | 1 | 0 | 0 | 0 | 0 | 0 | 0 | 0 | 1 | 1 | 1 | 0 |
| | TFF1 | 1 | 1 | 1 | 0 | 0 | 0 | 0 | 0 | 0 | 0 | 0 | 0 | 0 | 1 | 1 | 0 |
| | TM4SF20 | 0 | 0 | 1 | 0 | 1 | 0 | 0 | 0 | 0 | 0 | 0 | 0 | 0 | 0 | 1 | 0 |
| **Typical basal genes** | FAT2 | 0 | 0 | 0 | 0 | 0 | 0 | 0 | 1 | 1 | 0 | 0 | 0 | 0 | 0 | 0 | 1 |
| | KRT5 | 0 | 0 | 0 | 0 | 0 | 0 | 0 | 1 | 1 | 0 | 0 | 0 | 0 | 0 | 0 | 1 |
| | KRT6A | 0 | 0 | 0 | 0 | 0 | 0 | 1 | 1 | 1 | 0 | 0 | 0 | 0 | 1 | 0 | 1 |
| | KRT13 | 0 | 0 | 0 | 0 | 0 | 0 | 0 | 1 | 1 | 0 | 0 | 0 | 0 | 0 | 0 | 1 |
| | KRT14 | 0 | 0 | 0 | 0 | 0 | 1 | 0 | 1 | 1 | 0 | 0 | 0 | 0 | 1 | 0 | 1 |
| | RHCG | 0 | 0 | 0 | 0 | 0 | 0 | 0 | 1 | 1 | 0 | 0 | 0 | 0 | 0 | 0 | 1 |
| | TNNT3 | 0 | 0 | 0 | 0 | 0 | 0 | 0 | 0 | 1 | 0 | 0 | 0 | 0 | 0 | 0 | 1 |

| | Gene | Collisson_Classical | Moffitt_Classical | Bailey_Progenitor | Chan-Seng-Yue_Classical-A | Chan-Seng-Yue_Classical-B | Collisson_QM | Moffitt_Basal-like | Bailey_Squamous | Chan-Seng-Yue_Basal-A | Chan-Seng-Yue_Basal-B | Collisson_Exocrine-like | Bailey_ADEX | Bailey_Immunogenic | Puleo | Consensus_classical | Consensus_non-classical |
|---|---|---|---|---|---|---|---|---|---|---|---|---|---|---|---|---|---|
| **Typical acinar genes** | AMY2A | 0 | 0 | 0 | 0 | 0 | 0 | 0 | 0 | 0 | 0 | 0 | 1 | 0 | 0 | 0 | 1 |
| | CEL | 0 | 0 | 0 | 0 | 0 | 0 | 0 | 0 | 0 | 0 | 1 | 1 | 0 | 0 | 0 | 1 |
| | CELA3A | 0 | 0 | 0 | 0 | 0 | 0 | 0 | 0 | 0 | 0 | 1 | 1 | 0 | 0 | 0 | 1 |
| | CPA1 | 0 | 0 | 0 | 0 | 0 | 0 | 0 | 0 | 0 | 0 | 0 | 1 | 0 | 0 | 0 | 1 |
| | CPA2 | 0 | 0 | 0 | 0 | 0 | 0 | 0 | 0 | 0 | 0 | 0 | 1 | 0 | 0 | 0 | 1 |
| | CPB1 | 0 | 0 | 0 | 0 | 0 | 0 | 0 | 0 | 0 | 0 | 1 | 1 | 0 | 0 | 0 | 1 |
| | CTRB1 | 0 | 0 | 0 | 0 | 0 | 0 | 0 | 0 | 0 | 0 | 0 | 1 | 0 | 0 | 0 | 1 |
| | CTRB2 | 0 | 0 | 0 | 0 | 0 | 0 | 0 | 0 | 0 | 0 | 1 | 1 | 0 | 0 | 0 | 1 |
| | CTRC | 0 | 0 | 0 | 0 | 0 | 0 | 0 | 0 | 0 | 0 | 0 | 1 | 0 | 0 | 0 | 1 |
| | REG1B | 0 | 0 | 0 | 0 | 0 | 0 | 0 | 0 | 0 | 0 | 1 | 1 | 0 | 0 | 0 | 1 |
| **Typical immune genes** | IGKV1-12 | 0 | 0 | 0 | 0 | 0 | 0 | 0 | 0 | 0 | 0 | 0 | 0 | 1 | 0 | 0 | 1 |
| **Other genes present in Classical signature** | PRAP1 | 0 | 0 | 0 | 0 | 0 | 0 | 0 | 0 | 0 | 0 | 0 | 0 | 0 | 0 | 1 | 0 |
| | SLC13A2 | 0 | 0 | 0 | 0 | 0 | 0 | 0 | 0 | 0 | 0 | 0 | 0 | 0 | 0 | 1 | 0 |
| | B3GNT6 | 0 | 0 | 0 | 0 | 0 | 0 | 0 | 0 | 0 | 0 | 0 | 0 | 0 | 0 | 1 | 0 |
| | FEZF1 | 0 | 0 | 0 | 0 | 0 | 0 | 0 | 0 | 0 | 0 | 0 | 0 | 0 | 0 | 1 | 0 |
| **Other genes present in Non-classical signature** | DNER | 0 | 0 | 0 | 0 | 0 | 0 | 0 | 0 | 0 | 0 | 0 | 0 | 0 | 0 | 0 | 1 |
| | H19 | 0 | 0 | 0 | 0 | 0 | 0 | 0 | 0 | 0 | 0 | 0 | 0 | 0 | 0 | 0 | 1 |
| | TENM2 | 0 | 0 | 0 | 0 | 0 | 0 | 0 | 0 | 0 | 0 | 0 | 0 | 0 | 0 | 0 | 1 |

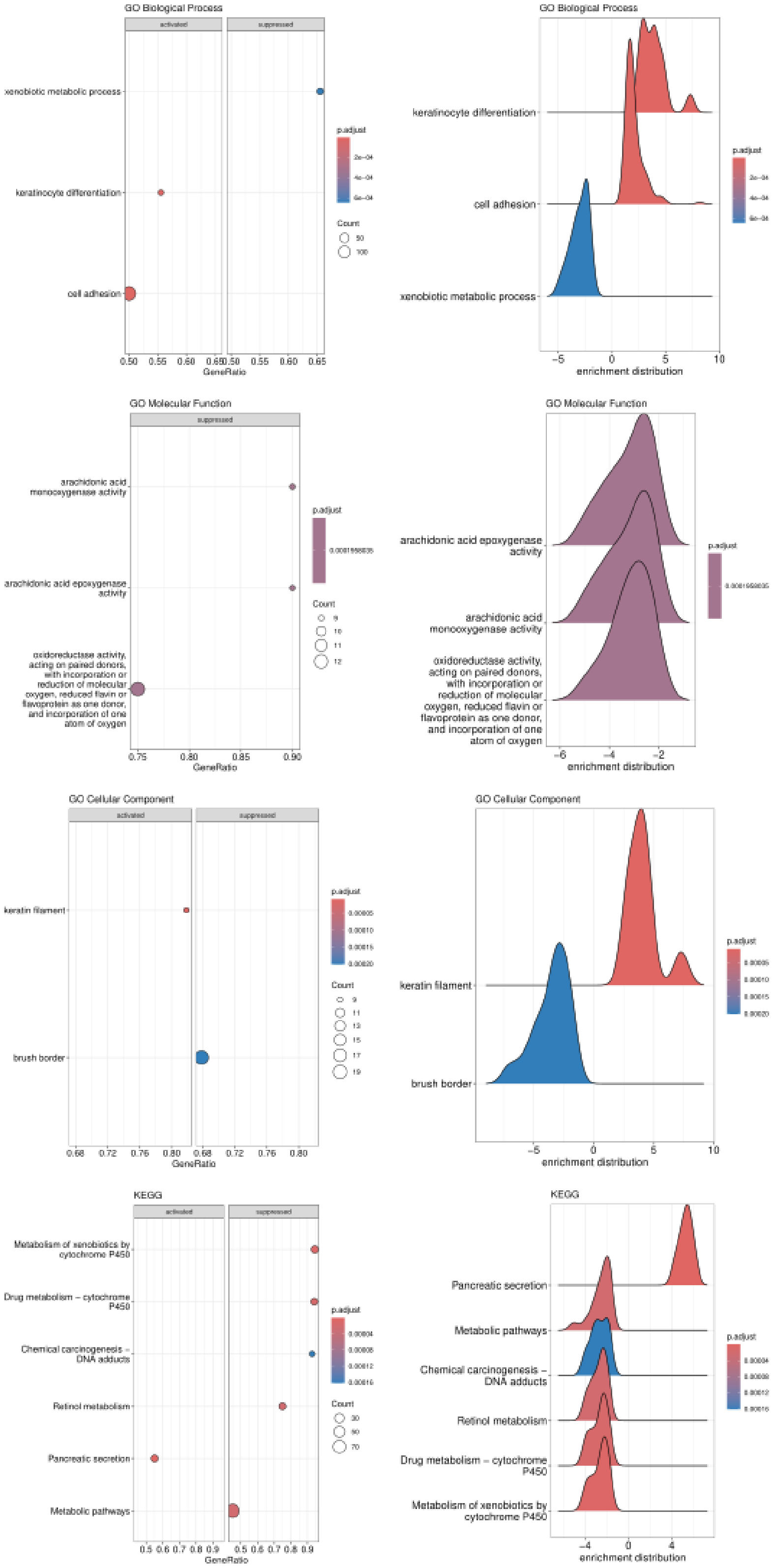


**Supplementary Figure 1.** Gene Set Enrichment Analysis of tumor consensus classes. *GeneRatio measures* the proportion of genes in a particular gene set that are found within the ranked lists of genes, while *enrichment distribution* quantifies the degree to which a gene set is overrepresented at the top or bottom of a ranked list of genes. GO, Gene Ontology; KEGG, Kyoto Encyclopedia of Genes and Genomes.

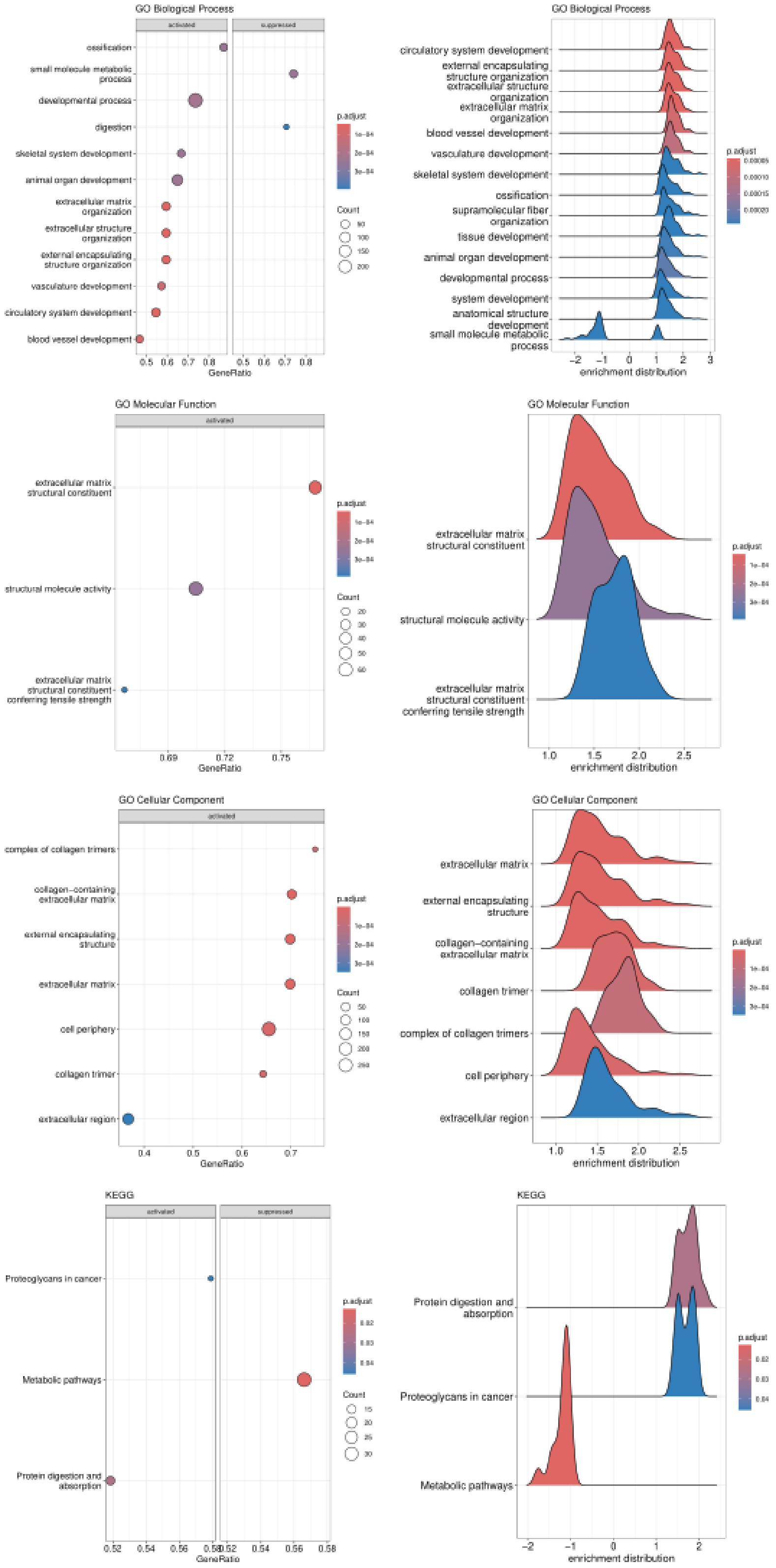


**Supplementary Figure 2.** Gene Set Enrichment Analysis of stroma consensus classes. Gene Ratio measures the proportion of genes in a specific gene set found within the ranked lists of genes, while enrichment distribution quantifies how much a gene set is overrepresented at the top or bottom of a ranked list. GO refers to Gene Ontology; KEGG stands for Kyoto Encyclopedia of Genes and Genomes.

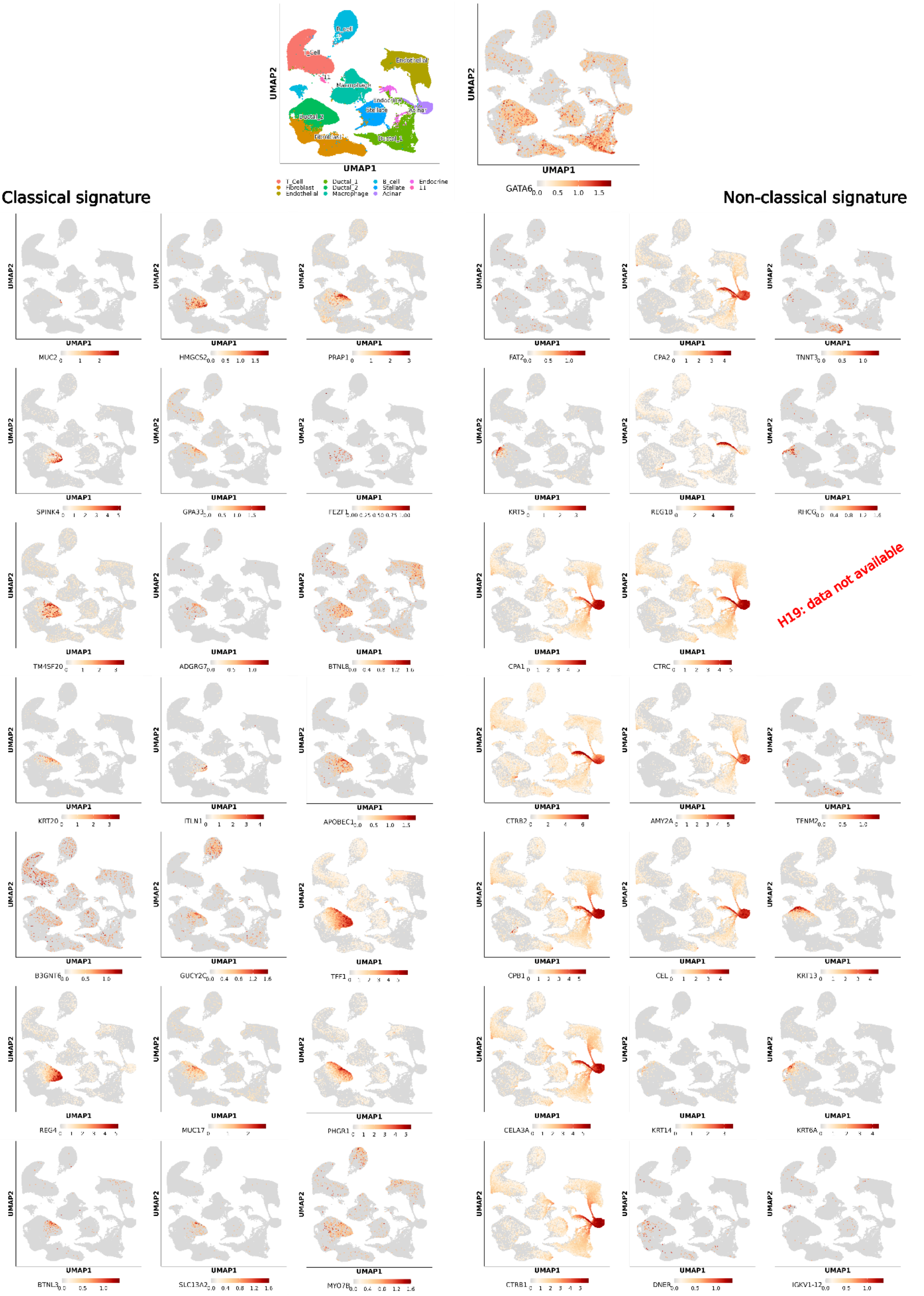


**Supplementary Figure 3.** Expression of genes associated with the Classical and Non-classical signatures in a single-cell dataset. At the top, the different clusters and the expression of GATA6 are shown for reference. Data for H19 was not included available.

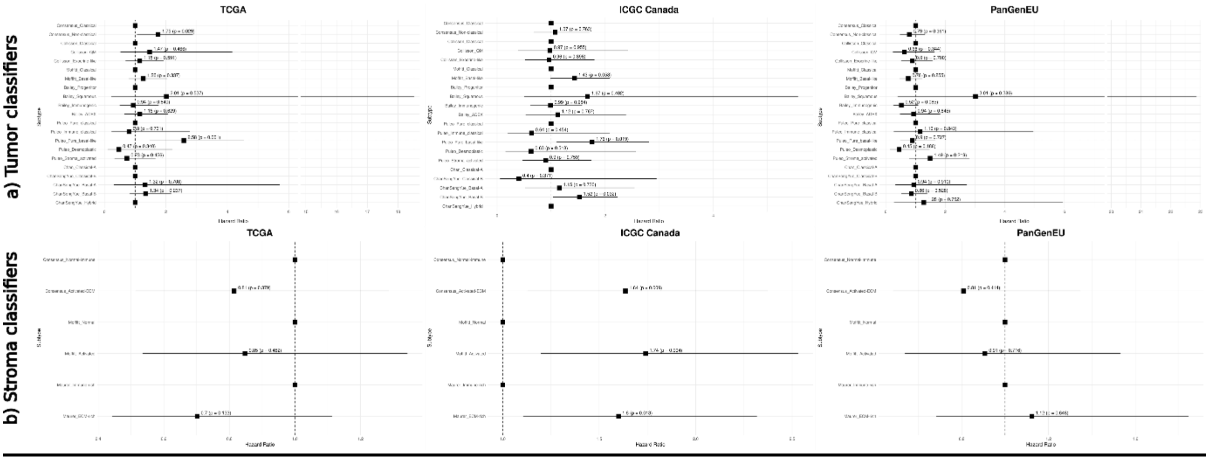


**Supplementary Figure 4.** Cox Proportional Hazards models for tumor and stroma classifiers in the training cohort. All models are adjusted for age, sex, and stage, and the hazard ratio and p-value are presented. ADEX (aberrantly differentiated endocrine-exocrine), ECM (extracellular matrix), ICGC (International Cancer Genome Consortium), QM (quasi-mesenchymal), TCGA (The Cancer Genome Atlas).
